# Simplexviruses successfully adapt to their host by fine-tuning immune responses

**DOI:** 10.1101/2020.07.28.226126

**Authors:** Alessandra Mozzi, Rachele Cagliani, Chiara Pontremoli, Diego Forni, Irma Saulle, Marina Saresella, Uberto Pozzoli, Mario Clerici, Mara Biasin, Manuela Sironi

## Abstract

Primate herpes simplex viruses are relatively harmless to their natural hosts, whereas cross-species transmission can result in severe disease. We performed a genome-wide scan for signals of adaptation of simplexviruses to hominins. We found evidence of positive selection in three glycoproteins, with selected sites located in antigenic determinants. Positively selected non-core proteins were involved in different immune-escape mechanisms. By expressing mutants of one of these proteins (ICP47), we show that the amino acid status at the positively selected sites is sufficient to induce HLA-G. HSV-1/HSV-2 ICP47 induced HLA-G when mutated to recapitulate residues in B virus, whereas the mutated version of B virus ICP47 failed to determine HLA-G expression. Thus, the evolution of ICP47 in HSV-1/HSV-2 determined the loss of an immunosuppressive effect, suggesting that simplexviruses tune immune responses to promote successful co-existence with their hosts. These results also help explain the high pathogenicity of B virus in humans.

## Introduction

Herpes simplex viruses (genus *Simplexvirus*, family *Herpesviridae*, order *Herpesvirales*), are dsDNA viruses that infect mammals, including humans and other primates. They have long genomes of approximately 155 kbp, organized into two regions of unique sequence (long (UL) and short (US)) flanked by direct or inverted repeats. The two unique sequences contain the great majority of the protein-coding regions, including core genes, which are shared among herpesviruses, and non-core genes, that are specific for members of the *Alphaherpesvirinae* subfamily and/or of the *Simplexvirus* genus only (McGeoch et al., 2006).

In analogy to other herpesviruses, the evolutionary history of simplexviruses was mainly characterized by coevolution and codivergence with their hosts (McGeoch et al., 2006). A known exception is represented by human herpes simplex virus 2 (HSV-2), which most likely originated from the cross-species transmission of an ancestor of chimpanzee herpesvirus 1 (PanHV-3) to an ancestor of modern humans, around 1.6 million years ago (Severini et al., 2013; Underdown et al., 2017; Wertheim et al., 2014). Thus, whereas most primates are infected by a single simplexvirus, humans host two: HSV-2 and human herpes simplex virus 1 (HSV-1). These viruses are present at high prevalence in human populations. Estimates vary with geography and reach 67% (HSV-1) and 11% (HSV-2) of the world population (Looker et al., 2015a; Looker et al., 2015b). Even if HSV-1 is primarily responsible for oro-facial lesions and HSV-2 for genital herpes (Arvin et al., 2007), both viruses can establish latency in trigeminal and lumbosacral ganglia, resulting in life-long infection (Arvin et al., 2007). Whereas a relatively low proportion of infected individuals show clinical manifestations during primary infection or reactivation (Tognarelli et al., 2019), simplexviruses can occasionally determine severe diseases such as infectious blindness, acute encephalitis, and neonatal invasive infection (Farooq and Shukla, 2012; Whitley, 2004).

In non-human primates (NHP), simplexvirus infections show symptoms and seroprevalence generally comparable to those of HSV-1 and HSV-2, and these viruses are species-specific in natural settings (Eberle and Jones-Engel, 2017). This feature, together with the near commensal relationship with their hosts, is in line with long-standing virus-host coevolution. Indeed, the consequences arising from disruption of the delicate balance established during millions of years of coexistence are evident when cross-species transmissions occur. For instance, macacine herpesvirus 1 (McHV1, also known as B virus) is almost asymptomatic in macaques, but infection of humans or African monkeys results in a severe, often fatal form of encephalomyelitis (Eberle and Jones-Engel, 2017; Loomis et al., 1981; Tischer and Osterrieder, 2010; Wilson et al., 1990).

Likewise, the transmission of HSV-1 from humans to marmosets or other New World monkeys is almost invariably fatal (Azab et al., 2018; Tischer and Osterrieder, 2010). These examples clearly testify how the viral and host genomes interact to determine the outcome of infection and highlight the potential zoonotic threat posed by simplexviruses. This also implies that simplexviruses must have adapted to their hosts and that the signatures of such adaptation may be detected using molecular evolution approaches. We thus performed a genome-wide scan of positive selection to identify variants in simplexvirus coding genes that arose during adaptation to the hominin lineage. As a proof of concept, we tested the functional effect of selected variants in *US12*, which encodes the ICP47 TAP inhibitor, a modulator of immune response.

## Results

### Selective patterns of catarrhini-infecting simplexvirus coding genes

We first explored the selective patterns of primate simplexvirus coding genes. We thus analyzed 6 complete genomes of simplexviruses that infect different primates, from hominins (HSV-1, HSV-2, and ChHV) to Old World African and Asian monkeys (CeHV-2, PaHV-2, and McHV-1) (Figure 1A, Table S1).

**Figure 1.**
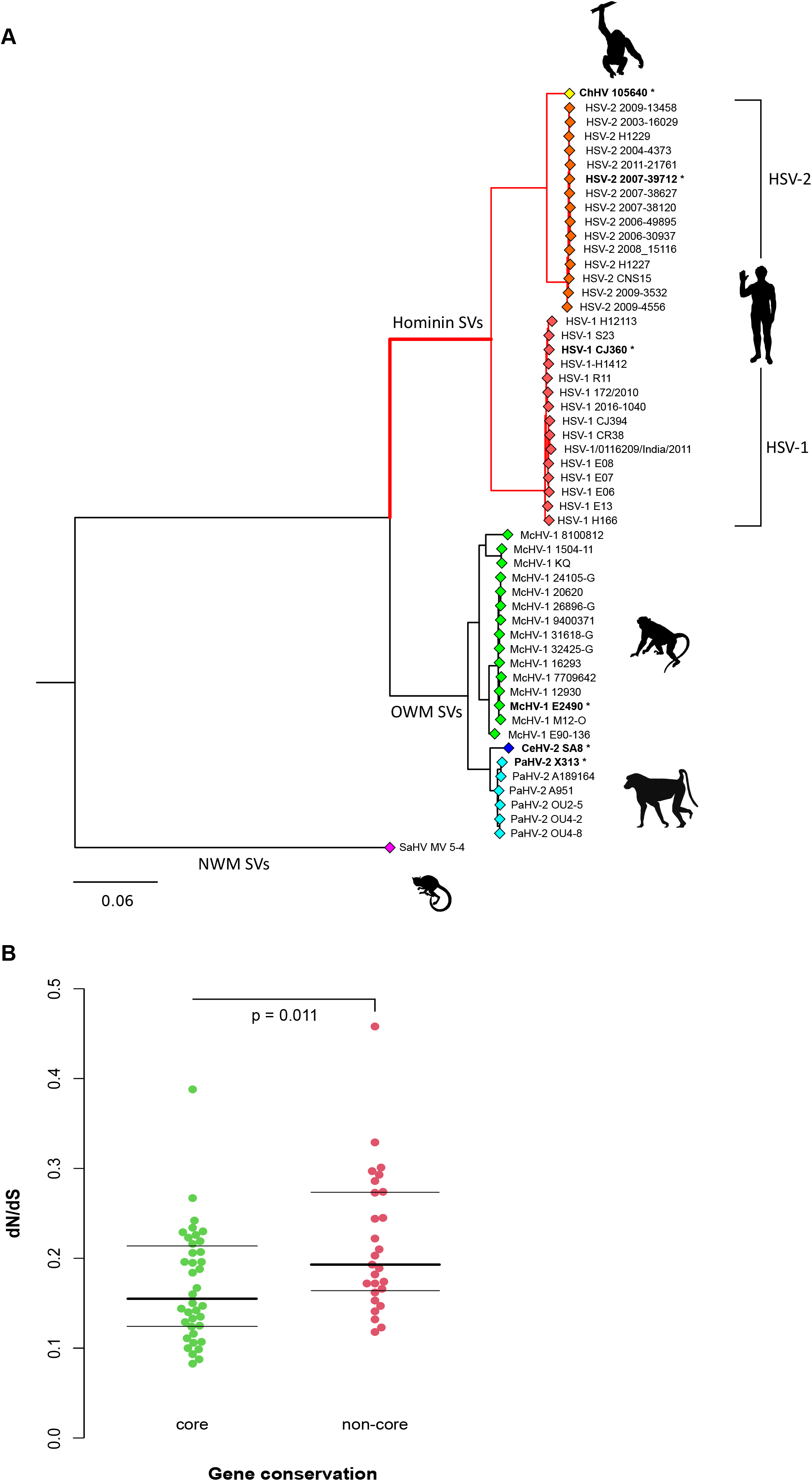
Selective patterns of primate simplexviruses. (**A**) A maximum-likelihood tree of glycoprotein B (encoded by the UL27 core gene) is drawn to exemplify the phylogenetic relationships among primate simplexviruses (Strain information and GeneBank IDs are reported in Table S1). The *Saimiriine alphaherpesvirus 1* (GeneBank ID: NC_014567) was used as the outgroup and the tree was constructed using PhyML (see methods). Asterisks denote viruses that were included in the analysis of selective patterns of catarrhini-infecting SVs. The hominin simplexvirus branch, that was specifically tested for episodic positive selection, is shown in red. (**B**) Comparison of dN/dS between *core* and *non-core* genes. The *p* value was calculated by the Wilcoxon Rank-Sum test.

Because high sequence diversity can affect evolutionary inference, viruses that infect New World primates were excluded from these analyses. Analysis of selective patterns was performed for all coding genes with reliable one-to-one orthologs. Gene sequences were rigorously filtered to ensure high quality alignments (see Materials and Methods). Genes for which few orthologous sequences were retrieved or with extended overlapping ORFs, were discarded (Table S2). The average non-synonymous substitution/synonymous substitution rate (dN/dS, also referred to as ω) was calculated for the resulting 65 genes (Table S3). Comparison of ω values among *core* genes (conserved among *Herpesviridae*, n=38) and *non-core* genes (specific to simplexviruses, n=27) indicated that these latter show lower evolutionary constraint (Wilcoxon Rank Sum test, p= 0.011) (Figure 1B, Table S3).

### Adaptive evolution in the hominin-infecting simplexvirus lineage

In order to assess whether adaptation to hominins drove the evolution of specific simplexvirus coding genes, we applied a branch-site test (Zhang et al., 2005) to an extended phylogeny of 53 viruses that infect hominins, Old World African monkeys, and Old world Asian monkeys (Figure 1A, Table S1). In the branch-site test, the branches of the tree are divided *a priori* into foreground and background lineages, and models that allow or disallow positive selection on the foreground lineage(s) are compared. The branch-site test can thus detect lineage-specific selected genes and sites (episodic positive selection). Herein, we set the branch leading to the hominin-infecting simplexviruses as foreground (Figure 1A).

After accounting for recombination (see Materials and Methods), we found evidence of adaptive evolution for 11 genes (16.9%). Positive selection in hominin simplexviruses similarly targeted *core* and *non-core* genes (selected fraction = 15.8% and 18.5%), irrespective of the higher selective constraint observed in *core* genes during viral evolution in catarrhini (Table S4).

We next analyzed positively selected sites. To be conservative, these were detected by the intersection of two approaches (see Materials and Methods). Among the *core* genes, we found evidence of episodic positive selection for three glycoproteins: gB (UL27), gH (UL22) and gM (UL10) (Figure 2 and Figure S1). *UL27* encodes the viral envelope glycoprotein B (gB), which is a major target antigen in herpesviruses (Malito et al., 2018). Both selected sites in gB are located in the ectodomain (Figure 2) and one of them, A334, is part of an epitope recognized by the SS55 neutralizing antibody (Cairns et al., 2014) (Figure 2). Interestingly, an R-to-Q substitution at residue 335, confers resistance to the SS55 mAb (Cairns et al., 2014). As for gH, two positively selected sites, Y85 and E170, flanked amino acids that, if mutagenized, confer resistance to the potent LP11 neutralizing antibody (Figure 2) (Chowdary et al., 2010). Because the LP11 antibody competes with gB for binding to the gH-gL complex, the gB binding site was proposed to be in close proximity to (or maybe overlapping) with the LP11 epitope surface (Chowdary et al., 2010). E170 is part of this surface, together with other sites we found under positive selection (Figure 2). Overall, these observations suggest that the selective pressure acting on these two glycoproteins is exerted by the host immune system.

**Figure 2.**
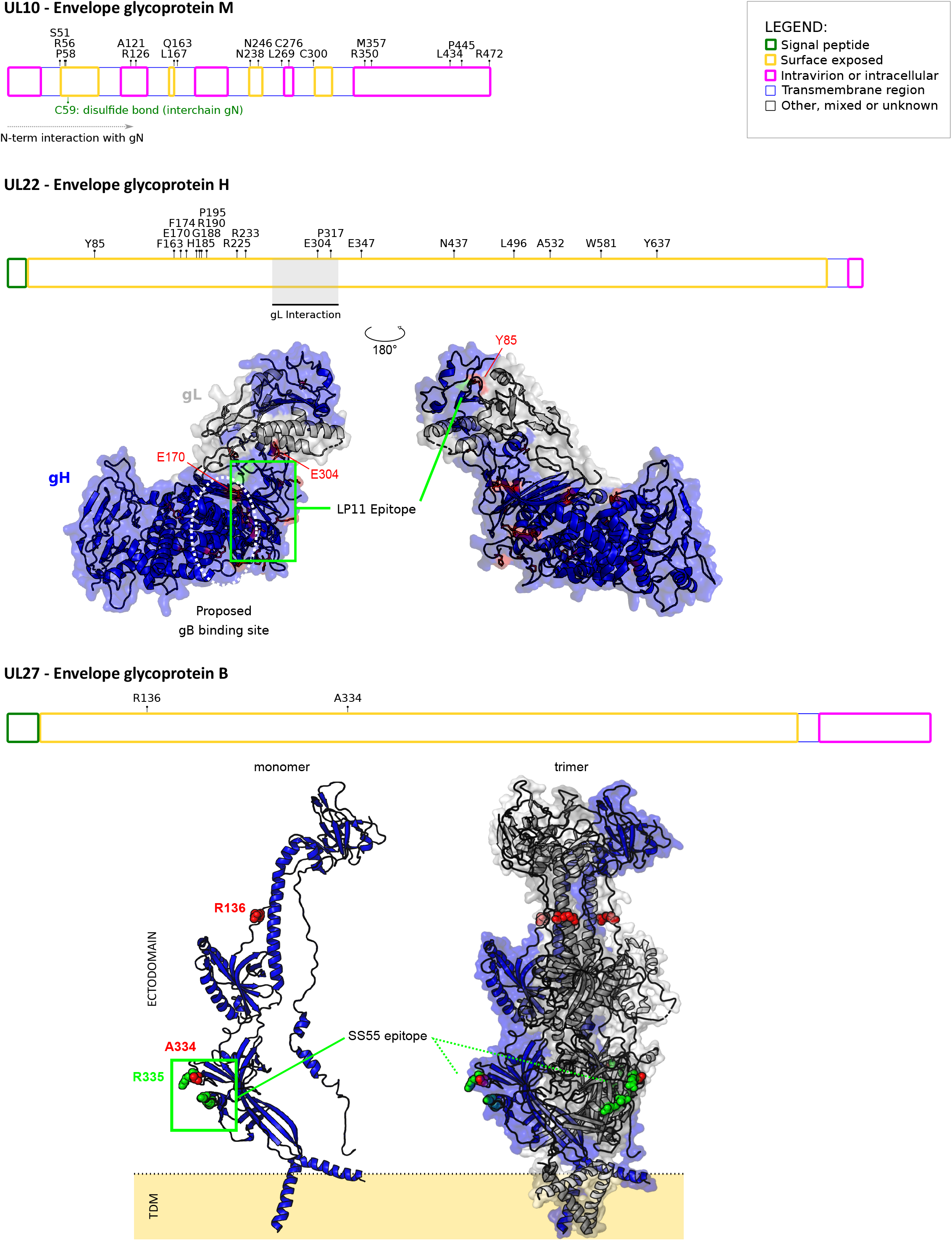
Positive selection in simplexvirus glycoproteins. Positively selected sites were mapped onto HSV-1 glycoproteins together with the location of functional domain/sites (grey). Topological features are color-coded according to the legend. For gH, positively selected sites (red) were mapped onto the three-dimensional structure of the gH-gL complex (blue and white, respectively; PDB ID: 3m1c) (Chowdary et al., 2010). The location of the LP11 epitope (green) and of the gB binding sites (white) are reported. The two views are rotated 180° around the vertical axis. For gB, positively selected sites were mapped onto the three-dimensional structures of the gB monomer (PDB ID: 6bm8) (Cooper et al., 2018) and the trimeric gB complex. This latter was obtained by a structural imposition of the monomer, using the 2gum structure as scaffold (Heldwein et al., 2006). The location of the SS55 epitope is reported in green. Positions refer to the reference HSV-1 strain 17 (NC_001806).

We also found many positively selected sites in gM (UL10); the N-terminus of gM is predicted to interact with the glycoprotein N (gN), to form a stable complex, which modulates the viral fusion machinery (El Kasmi and Lippé, 2015). Three of the positively selected sites (S51, R56, P58) we found, are located in the surface exposed region of gM, just upstream the cystein (C59) residue which is responsible for an interchain disulphide bond that stabilize the gM-gN complex (Striebinger et al., 2016), strongly suggesting that these residues could contribute to gM-gN interaction. Several other positively selected sites were located along the whole sequence of gM (Figure 2).

Among *non-core* genes showing evidence of positive selection, four (*UL46, US8, US1*, and *US12*) are involved in different immune-escape mechanisms. US8 codes for glycoprotein E (gE) that, in complex with gI, forms an Fc receptor for immunoglobulin G (IgG) (Dubin et al., 1991; Sprague et al., 2006). The gE-gI complex binds the Fc region of IgG leading to an antibody bipolar bridging on infected cells, preventing IgG-mediated immune response. In US8, six positively selected sites were found in the protein domain involved in Fc interaction; among them, E227 and G313 lies at Fc interaction surface boundaries (Figure 3) (Sprague et al., 2006).

**Figure 3.**
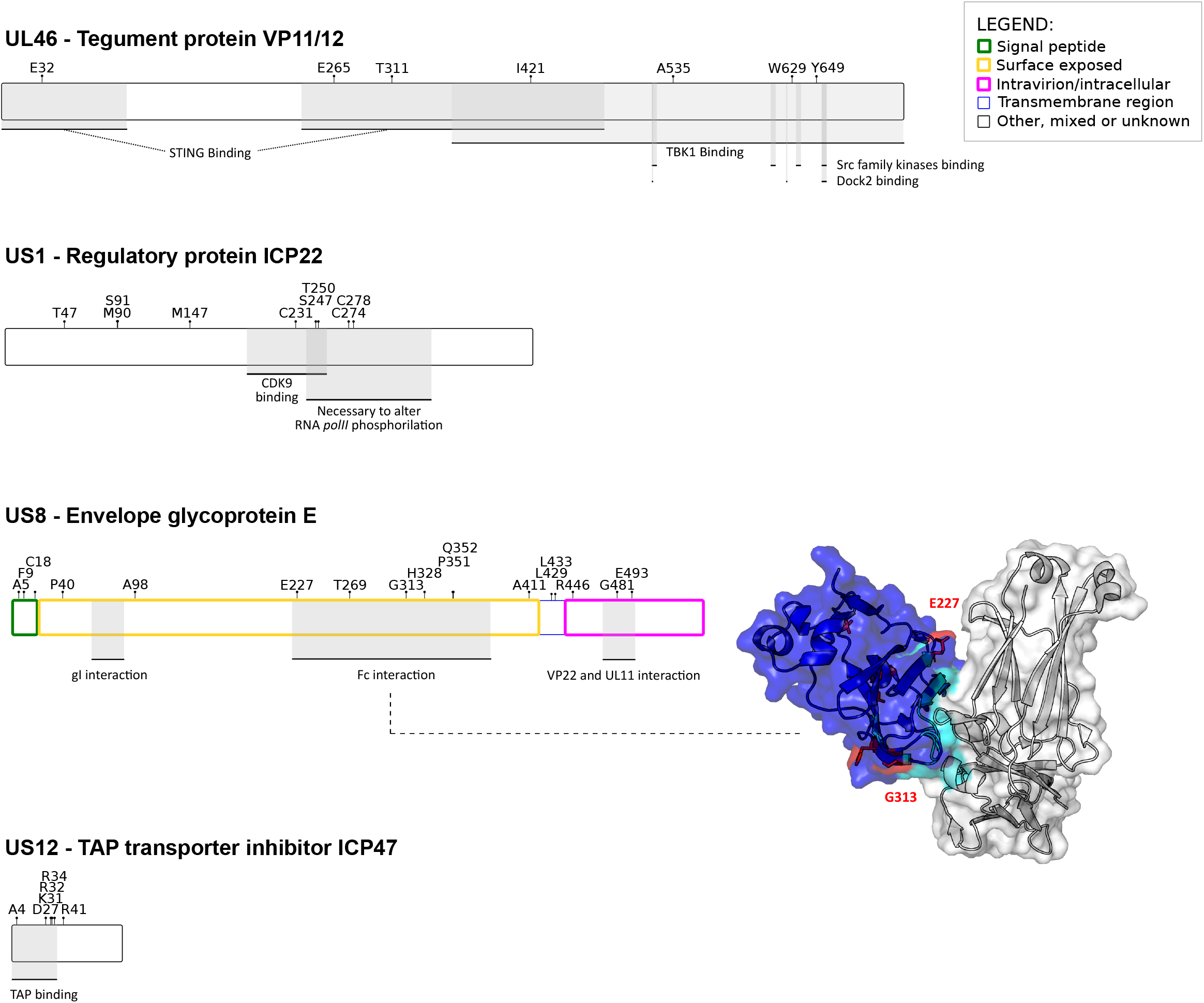
Positive selection in SVs proteins involved in host immune system-escape. Proteins and positively selected sites are reported as in Figure 2. For gE, the three-dimensional structure of the complex with Fc (PDB ID:2gj7) is reported (Sprague et al., 2006). gE is represented in blue, with the Fc interaction surface in cyan. Positively selected sites are in red. Positions refer to the reference HSV-1 strain 17 (NC_001806).

UL46 encodes an abundant tegument protein that mediates viral evasion from foreign DNA-sensing pathways (Deschamps and Kalamvoki, 2017). In particular, the UL46 protein of HSV-1 interacts with both TMEM173/STING and TBK1 through separate domains and blocks the DNA-sensing pathway. We detected positively selected sites both in the STING and in the TBK1 binding regions (Figure 3). US1 encodes the ICP22 protein (Figure 3), a general transcription regulator that also down-modulates the expression of CD80 in dendritic cells (Matundan and Ghiasi, 2019). Finally, US12 encodes the ICP47 protein, which down-regulates the expression of major histocompatibility complex (MHC) class I molecules on the cell surface (Früh et al., 1995; Hill et al., 1995). In particular, the ICP47 proteins of HSV-1 and HSV-2 act as inhibitors of the transporter associated with antigen processing (TAP), which translocates antigenic peptides into the endoplasmic reticulum lumen for loading onto MHC class I (HLA-ABC) molecules (Früh et al., 1995; Hill et al., 1995; Tomazin et al., 1998). The TAP-biding region resides in the N-terminal portion of the ICP47 protein, where all the positively selected sites are located (Galocha et al., 1997; Matschulla et al., 2017) (Figures 3 and 4A-B).

**Figure 4.**
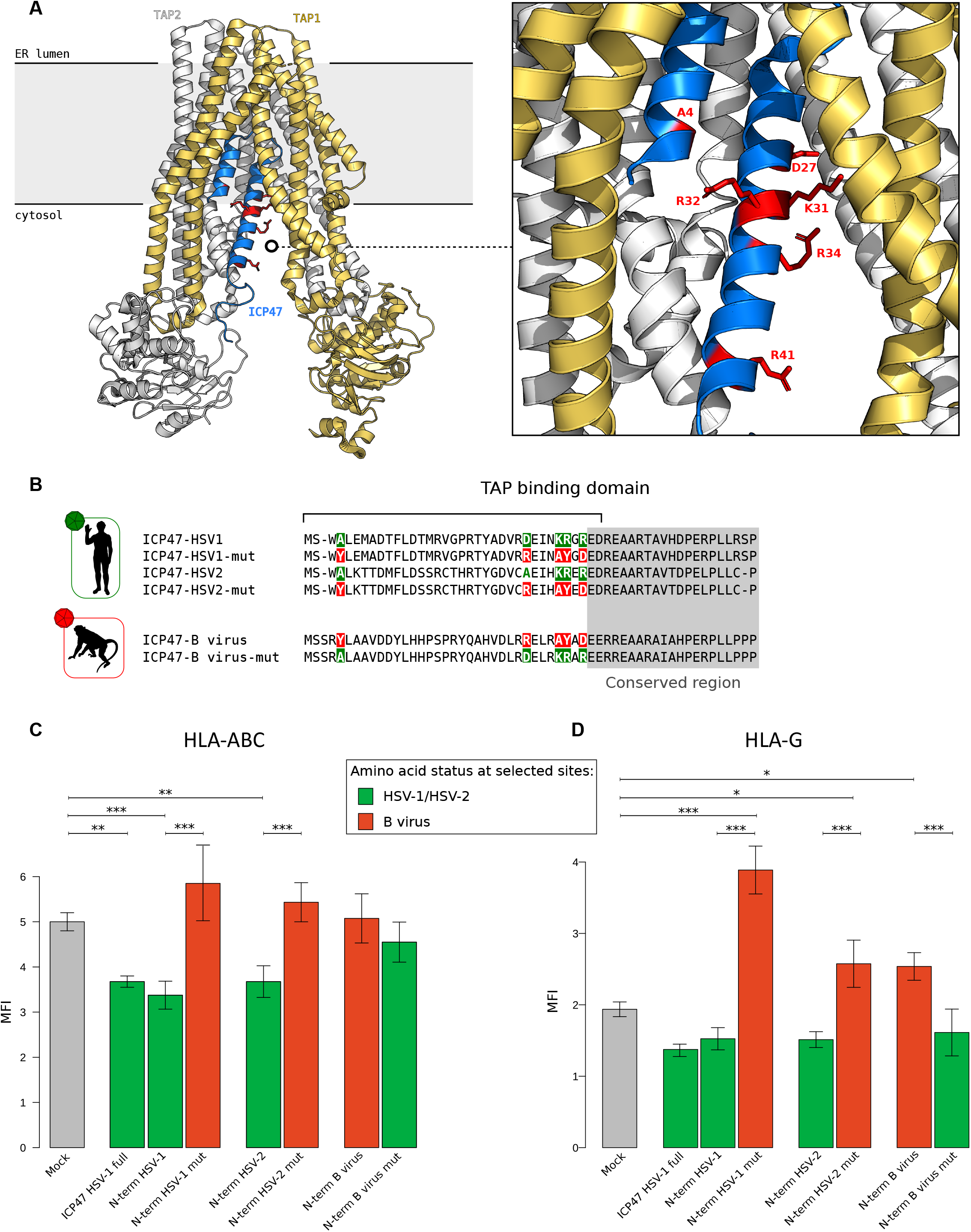
Functional characterization of positive selected sites of US12 (ICP47). (**A**) Ribbon representation of the three-dimensional structure of ICP47 bound to the TAP transporter, which in turn is composed by two subunits, TAP1 (light orange) and TAP2 (white) (PDB ID: 5u1d) (Oldham et al., 2016). Positively selected sites are represented as red sticks in the enlargement. (**B**) Schematic view recapitulating the amino state of positively selected sites tested in our analyses. HLA-ABC (**C**) and HLA-G (**D**) expression at the cell surface. Jurkat cells were transfected with the ICP47 constructs and the amounts of total HLA-ABC or HLA-G antigen was quantified by cytofluorimetry after 48 hours. MFI (mean fluorescence intensity) bar plots represent the mean and standard deviation of four replicates. *p* values were calculated using Tukey post hoc tests.

### Positively selected sites in *US12* modulate the surface expression of MHC class I molecules

The N-terminal region of ICP47 is poorly conserved across hominin-infecting SVs, and considerable divergence is also observed between HSV-1 and HSV-2, which however bind and inhibit human TAP (Tomazin et al., 1998). In particular, the 55 N-terminal residues of HSV-1 ICP47 are sufficient to interact with and inhibit TAP (Galocha et al., 1997; Matschulla et al., 2017). Conversely, a previous study indicated that the ICP47 protein encoded by B virus lacks the ability of HLA-ABC down-regulation, although up-regulation of HLA-E and HLA-G was observed during infection (Vasireddi and Hilliard, 2012). We thus investigated whether the positively selected sites in ICP47 modulate the different ability of simplexvirus proteins to regulate HLA-ABC expression, and if the ICP47 protein of B virus is responsible for HLA-G up-regulation. To this aim, we designed constructs carrying the TAP binding domains of HSV-1, HSV-2, or the corresponding region of B virus ICP47. Two additional constructs carried the HSV-1 or HSV-2 ICP47 N-terminal domain mutagenized at the positively selected sites to recapitulate the amino acid state observed in the macaque virus. In turn, mutations reproducing the amino acids observed in the human virus were introduced in the B virus N-terminal domain (Figure 4B).

These constructs, together with a plasmid expressing the full-length HSV-1 ICP47 protein, were transiently transfected in Jurkat cells and the surface expression of HLA-ABC molecules was evaluated by cytofluorimetry. As expected, a significant effect of the plasmids on HLA-ABC expression was evident (ANOVA, F = 19.55, P = 7.22 x 10^−8^), whereas the differences among replicates were not significant (F = 2.57, P = 0.08). Tukey post hoc tests indicated that the full-length ICP47 protein and the two TAP binding domains of HSV-1 and HSV-2 significantly reduced HLA-ABC expression compared to mock transfected cells (Figure 4C). No difference was observed between the two short constructs and the complete ICP47 protein. Mutation of the positively selected sites in both the HSV-1 and HSV-2 TAP biding domains of ICP47 totally abolished these effects (Figure 4C), suggesting that the selected sites play an important role in TAP binding. In line with previous results (Vasireddi and Hilliard, 2012), the ICP47 domain of B virus did not affect HLA-ABC expression, However, mutation of the selected sites to recapitulate the amino acids observed in the HSV-1/HSV-2 molecules was not sufficient to restore TAP inhibition (Figure 4C). Overall, these results indicate that the positively selected sites are not the sole determinants of TAP binding. We next assessed the effect of the different ICP47 constructs on HLA-G expression. Again, a significant effect for the constructs (ANOVA, F = 53.28, P = 5.91 x 10^−12^) but not for the replicates (F = 0.73, P = 0.55) was observed. The short N-terminal domain of B virus ICP47 was sufficient to significantly increase HLA-G expression compared to mock-transfected cells (Figure 4D). Mutation of the positively selected sites to those observed in HSV-1 and HSV-2 fully abrogated the increased expression of HLA-G. Interestingly, whereas the expression of full or partial HSV-1/HSV-2 ICP47 did not affect HLA-G expression, introduction of mutations that recapitulate amino acids observed in B virus conferred to both ICP47 N-terminal domains the ability to induce HLA-G (Figure 4D). These results indicate that the modulatory effect of B virus on HLA-G expression is mediated by the N-terminal domain of ICP47 and that the positively selected sites are the major determinants of HLA-G regulation. Clearly, the effect on HLA-G expression must be TAP-independent.

## Discussion

Primate simplexviruses are often regarded as an epitome of virus-host coevolution and codivergence (Eberle and Jones-Engel, 2017; McGeoch et al., 2006). These viruses establish life-long infections and usually cause little harm to their hosts, whereas periodic viral reactivation allows transmission in the population. Indeed, virulence and host range are often interconnected traits in viruses (Rothenburg and Brennan, 2020), which are expected to evolve to maximize their transmission potential in the host and to tune their virulence accordingly.

Whereas several herpesviruses are unable to infect species other than their natural host, the occasional cross-species transmission of primate simplexviruses has been documented several times, indicating that few barriers exist in terms of infection potential (Azab et al., 2018). However, most spill-overs result in a very severe disease in the new host, especially when the phylogenetic distance from the original host is considerable (Azab et al., 2018). For instance, HSV-1 infection is almost invariably fatal in New World monkeys, whereas limited data on gorillas and Old World monkeys suggest that the symptoms are milder (Gilardi et al., 2014; Tischer and Osterrieder, 2010). The best known example of the severe effects of cross-species transmission is that of B virus. Although the virus is rarely acquired, even in people who are in frequent contact with macaques, mortality due to central nervous system involvement is extremely high when infection occurs (Azab et al., 2018; Eberle and Jones-Engel, 2018; Tischer and Osterrieder, 2010).

These observations clearly indicate that simplexviruses have been adapting to their hosts to balance virulence and transmission. Such a balance is most likely the result of multiple interactions between virus- and host-encoded factors, and the interplay between the host immune response and the viral evasion strategies is expected to determine the outcome of infection. We thus searched for signals of adaptation of simplexviruses to their hominin hosts. Specifically, we applied a branch-site test, which is well-suited to identify episodic positive selection - i.e., selection events that occurred on a specific branch of a phylogeny. Among *core* genes, we found evidence of episodic positive selection in three glycoproteins, namely gB, gM, and gH, all of which contribute to virus cell entry via membrane fusion (Arii and Kawaguchi, 2018; El Kasmi and Lippé, 2015). For gB and gH we found that some of the positively selected sites map to antigenic determinants, suggesting that the host adaptive immune response represents the underlying selective pressure. Moreover, these glycoproteins participate in other processes that contribute to the alteration of the host immune responses. In fact, gB affects the trafficking of MHC class II molecules and diverts them to the exosome pathway (Temme et al., 2010), whereas gH interacts with both αvβ3-integrin and TLR2, which sense the virus and activate the innate immune response (Gianni et al., 2012; Leoni et al., 2012). Both gH and gM were also reported to counteract tetherin, a cellular restriction factor for several enveloped viruses(Blondeau et al., 2013; Liu et al., 2015). In line with the view that hosts and viruses are engaged in genetic conflicts, tetherin was shown to have evolved under positive selection in primates (Gupta et al., 2009; Lim et al., 2010; McNatt et al., 2009). Indeed, this is a general finding for a number of genes involved in defense mechanisms, which display unusually rapid rates of evolution in response to the selective pressure imposed by pathogens (Sironi et al., 2015). Clearly, several infectious agents can insist on the same defense pathway, implying that pathogens are faced with a fast-evolving array of host defense mechanisms. For instance, STING, a stimulator of interferon responsive genes, which is positively selected in primates (Mozzi et al., 2015), is targeted by several viruses. We found three positively selected sites in the STING-binding domain of UL46, suggesting virus adaptation to modulate interaction with the host molecule. Another cellular system commonly antagonized by viruses is the antigen processing and presentation pathway, many components of which show rapid evolutionary rates (Forni et al., 2014). In particular, different herpesviruses employ distinct strategies to interfere with the antigen presentation pathway, thus protecting themselves from the host immune response (van de Weijer et al., 2015; Verweij et al., 2015). In addition to the above-mentioned effect of gB on MHC class II sorting, simplexviruses express the ICP22 protein, which is positively selected and down-modulates CD80 (Matundan and Ghiasi, 2019), as well as the ICP34.5 protein (the product of *RL1*). ICP34.5, a neurovirulence factor that blocks MHC II expression on the surface of infected cells (Trgovcich et al., 2002). Due to the small number of confirmed orthologs of *RL1* we could no test whether positive selection acted on this gene.

We instead analyzed the selection pattern of *US12*, which encodes ICP47. All but one of the positively selected sites we detected were located within the N-terminal domain. For the HSV-1 ICP47 protein, this region is sufficient to bind TAP and freeze it in an inactive conformation (Galocha et al., 1997; Matschulla et al., 2017). Because peptide loading is necessary to allow folding of HLA class I molecules in their active configuration, this in turn results in the retention of HLA-ABC molecules in the endoplasmic reticulum. HLA-ABC down-regulation prevents the recognition of infected cells by CD8^+^ T-lymphocytes, which explains why TAP inhibition is a common viral strategy of immune subversion (Hill et al., 1995). The TAP binding activity of ICP47 was demonstrated for both the HSV-1 and HSV-2 proteins, although sequence similarity is limited in the N-terminal portion. Conversely, infection with B virus does not result in the down-modulation of HLA-ABC expression (Vasireddi and Hilliard, 2012). We thus reasoned that the selected sites might underlie the different ability of simplex viruses to inhibit TAP. However, our data indicate that, although the amino acid status at these sites is clearly important, as their mutation in HSV-1/HSV-2 ICP47 restored HLA-ABC expression to the same level as non-transfected cells, they do not represent the sole determinants of TAP binding. In fact, when the amino acids observed in the human viruses were introduced in the N-terminus of B virus ICP47, no HLA-ABC down-modulation was observed. Conversely, the amino acid status at the positively selected sites is sufficient to determine HLA-G up-regulation. In fact, the N-terminal domains of both HSV-1 and HSV-2 ICP47 induced HLA-G when mutated to recapitulate residues in B virus. Conversely, the mutated version of B virus ICP47 failed to determine HLA-G expression. Overall, these results imply that the ability of B virus to induce HLA-G resides in the N-terminal domain of ICP47 and that it does not depend on TAP. This is consistent with the notion that HLA-G can be loaded with peptides by both TAP-dependent and TAP-independent pathways (Lee et al., 1995). The mechanism underlying the up-regulation of HLA-G by B virus ICP47 remains unexplored, and further experiments will thus be required to determine how the positively selected sites exert their effect. As a corollary, our data indicate that the short region of ICP47 we analyzed herein could be used as an inducer of HLA-G expression, which is regarded as a potential biotherapy in allogenic transplantation (Deschaseaux et al., 2011).

The reason why related viruses use the same protein to differentially modulate host responses remains to be clarified. The loss of TAP-binding activity by B virus ICP47 may represent a strategy to limit NK cell activation (Vasireddi and Hilliard, 2012). In fact, reduced HLA-ABC expression on the cell surface results in NK-mediated killing, unless inhibitory ligands are also expressed (Früh et al., 1995; Hill et al., 1995; Huard and Früh, 2000). Indeed, NK cells play a central role in limiting HSV-1/HSV-2 infection, as demonstrated by mouse models (Rager-Zisman et al., 1987), as well as by the extremely severe infection outcome in humans with genetic defects resulting in low/absent NK cell counts (Orange, 2013). It was instead suggested that B virus, due to its lack of TAP-inhibitory activity, does not trigger NK responses(Vasireddi and Hilliard, 2012). In addition, at least in human cells, this virus up-regulates HLA-G (Vasireddi and Hilliard, 2012), which is associated with diverse immunosuppressive functions, including inhibition of T cell and NK cell responses (Morandi et al., 2016). On one hand these observations might account for the extreme virulence of B virus in humans. On the other, as noted elsewhere(Eberle and Jones-Engel, 2018), they do not explain why infection in macaques is poorly pathogenic. Notably, though, rhesus macaques do not express the ortholog of HLA-G, as it is a pseudogene (Boyson et al., 1997). Through alternative splicing, the Mamu-AG gene of these non-human primates encodes glycoproteins functionally similar to HLA-G (Slukvin et al., 2000), which is also alternatively spliced. Mamu-AG shares several features with human HLA-G, including a role in the establishment of maternal-fetal immune tolerance, but it is phylogenetically more similar to HLA-A (Boyson et al., 1997). It is thus possible that Mamu-AG glycoproteins are not up-regulated by ICP47 and that, therefore, infection in macaques elicits weaker immunomodulatory effects, eventually resulting in mild presentation. Addressing this point will require further analyses and the generation of antibodies against Mamu-AG, which are not commercially available.

In summary, we performed a genome-wide scan of positive selection on the hominin simplexvirus branch. We detected several positively selected sites, many of which most likely evolved in response to immune-mediated selective pressure. As these sites were positively selected, they are expected to affect some viral traits, as phenotypes are the ultimate target of selection. As a proof of concept, we tested the functional effects of positively selected sites in ICP47. Such sites were found to be sufficient to determine the inability of the viral protein to up-regulate HLA-G expression. Thus, the evolution of ICP47 in HSV-1/HSV-2 determined the loss of an immunosuppressive effect, suggesting that the trait under selection was decreased virulence. This possibility parallels findings in human cytomegalovirus, another herpesvirus, whereby different mechanisms promoting viral temperance were described (Dunn et al., 2003; Mozzi et al., 2020). These analyses may also suggest that closely related viruses finely tune the balance between immunosuppressive and immunostimulatory pathways to promote successful co-existence with their primate hosts.

## Materials and Methods

### Sequences and alignments

Viral genome sequences were retrieved from the NCBI (http://www.ncbi.nlm.nih.gov/) database. A detailed list of accession number is reported in Table S1. Alignments of whole genome sequences were performed with Progressive MAUVE 2.3.1, using default parameters (Darling et al., 2004; Darling et al., 2010). For each viral genome, we retrieved coding sequences of all annotated ORFs. Orthology was inferred according to MAUVE attribution and to genome annotation.

Gene alignments were generated using MAFFT (Katoh and Standley, 2013), setting sequence type as codons. Unreliably aligned codons were filtered using GUIDANCE2 (Sela et al., 2015) with a cutoff of 0.90 (Privman et al., 2012). The resulting alignments were manually inspected. Only reliable one-to-one orthologs were included in the subsequent analyses (Table S3).

### Selective patterns in primate-infecting simplexviruses

The average dN/dS parameter was calculated using the single-likelihood ancestor counting (SLAC) method (Kosakovsky Pond and Frost, 2005), using a phylogeny of 6 SVs infecting different primate species (Table S1).

Phylogenetic trees were generated with the phyML program (version 3.1), by applying a General Time Reversible (GTR) model plus gamma-distributed rates and 4 substitution rate categories, a fixed proportion of invariable sites, and a BioNJ starting tree (Guindon et al., 2009).

Differences in dN/dS among catarrhini-infecting SVs genes grouped on the basis of gene conservation in the *Herpesvirales* order (Davison, 2007) were evaluated using the Wilcoxon rank sum test.

### Detection of positive selection in the hominin-infecting simplexvirus lineage

We analyzed a viral phylogeny composed of 53 catarrhini-infecting viral strains of *Simplexvirus* genus. Specifically, we include 22 fully-sequenced strains infecting Old word monkey species (i.e., macaques and baboons), 1 strain infecting chimpanzee, and 30 strains infecting humans (both HSV-1 and HSV-2, n=15 respectively). HSV-1 and HSV-2 strains were selected from clinical isolates with no history of passaging in cell culture, sampled in different countries in order to have an heterogeneous pool of viral genomes representative of the diversity among circulating strains (Table S1).

Analyses were performed on the same phylogeny of catarrhini-infecting simplexviruses (see above); for each coding-gene, phylogenetic trees were reconstructed using phyML. Each alignment was screened for the presence of recombination using GARD (Kosakovsky Pond et al., 2006), a genetic algorithm implemented in the HYPHY suite (version 2.2.4). When evidence of recombination was detected (*p* value<0.01), the coding alignment was split accordingly; sub-regions were than used as the input for molecular evolution analyses. Only resulting alignments that, after GUIDANCE filtering had a length ≥ 250 nt were considered for subsequent analyses. Episodic positive selection on the Hominin-infecting simplexviruses branch was detected by applying the branch-site likelihood ratio tests from codeml (“test 2”) (Zhang et al., 2005). In this test, a likelihood ratio test is applied to compare a model (MA) that allows positive selection on the foreground lineages with a model (MA1) that does not allow such positive selection. Twice the difference of likelihood for the two models (ΔlnL) is then compared to a χ^2^ distribution with one degree of freedom (Zhang et al., 2005). The analyses were performed using an F3X4 codon frequency models. An FDR correction was applied to account for multiple tests.

To identify sites evolving under positive selection, we used BEB analysis from MA (with a cutoff of 0.90) and the Mixed Effects Model of Evolution (Murrell et al., 2012) (MEME, cutoff of 0.1), that allows ω to vary from site to site and from branch to branch at a site. To limit false positives, only sites confirmed by both methods were considered as positively selected.

### Plasmids

The coding sequences of ICP47 N-terminus from HSV-1 (55aa, YP_009137148), HSV-2 (55aa, YP_009137225.1), and B-virus (56aa, NP_851932) were synthesized and cloned in pCMV6-Entry vector by Origene custom service. The pCMV6 vectors coding for the corresponding mutagenized sequences were synthesized and cloned as well (Figure 4B).

### Cell culture and transfection

Jurkat cells were cultured in RPMI complete media without antibiotics and supplemented with 10% Fetal Bovine Serum (FBS). Cells were cultured at 37 °C and 5% CO_2_ in Forma Steri-Cycle CO_2_ incubator (Thermo). Every 3 days, cells were split to 0.5–1 × 10^6^ cells/ml in a T25 culture flask with fresh media. ~5×10^5^ Jurkat cells were electroporated in a solution of R-buffer (100μL; Invitrogen) containing 1ug of plasmid (HSV-1 full, N-term HSV-1, N-term HSV-1-mut N-term HSV-2, N-term HSV-2 mut N-term B virus, N-term B virus mut) using a Neon^®^ Transfection System (Invitrogen) under the recommended electroporation condition (1350 V, 10 ms, 3 pulse). The transfected cells were then seeded into 24-well plate. All experiments were run in four replicates and cells electroporated without plasmid were considered as the control (mock).

Post transfection Jurkat cell viability was ≥90% as determined by an automatic cell counter (Digital Bio, NanoEnTek Inc, Korea).

### Immunofluorescent staining and Flowcytometry analysis

PBMCs were stained with HLA-ABC PE (Clone W6/32, eBioscience), and HLA-G PE-Cy7 (Isotype IgG2 Mouse, Clone 87G, eBioscience), for 15 min at room temperature in the dark. After incubation, Jurkat cells were washed and resuspended in PBS.

Flow cytometric analyses were performed after 2 days post-transfection using a Beckman Coulter Gallios Flow Cytometer equipped with two lasers operating at 488 and 638 nm, respectively, interfaced with Gallios software and analyzed with Kaluza v 1.2. Two-hundred-thousand events were acquired and gated on HLA-ABC or HLA-G for Jurkat cells.

Data were collected using linear amplifiers for forward and side scatter and logarithmic amplifiers for fluorescence (FL)1, FL2, FL3, FL4, and FL5. Samples were first run using isotype control or single fluorochrome-stained preparations for color compensation. Rainbow Calibration Particles (Spherotec, Inc. Lake Forest, IL) were used to standardize flow-cytometry results.

Results were expressed as Mean Intensity Fluorescence (MFI) of HLA-ABC and HLA-G on Jurkat cells.

## Funding

This work was supported by the Italian Ministry of Health (“Ricerca Corrente 2019-2020” to MS, “Ricerca Corrente 2018-2020” to DF)

## Competing interests

The authors declare no conflict of interest

## Notes

### Competing Interest Statement

The authors have declared no competing interest.

