## Supplementary figure and tables for "Simplexviruses successfully adapt to their host by fine-tuning immune responses"

**Supplementary Figures:**

**Figure S1. Positive selection in UL26, UL29, UL36 and UL55.** Positively selected sites and functional domains were mapped onto HSV-1 proteins, as in Figure 2 and 3. For UL36, given the extended length of the protein sequence, positively selected sites were reported in the enlargement below.

**Supplementary Tables:**

**Supplementary Table S1. List of viral genome sequences**

**Supplementary Table S2. Herpes simplex genes excluded from the branch-sites analysis**

**Supplementary Table S3. List of analyzed genes and dN/dS values**

**Supplementary Table S4. Likelihood ratio test (LRT) statistics for models of variable selective pressure on the Hominin-infecting SVs branch.**

### UL26 - Capsid scaffolding protein

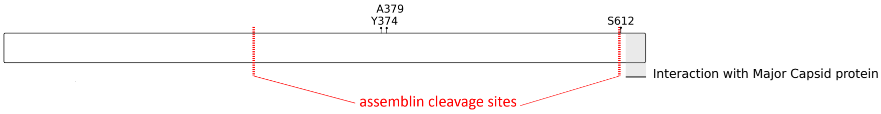


### UL29 - Major DNA-binding protein

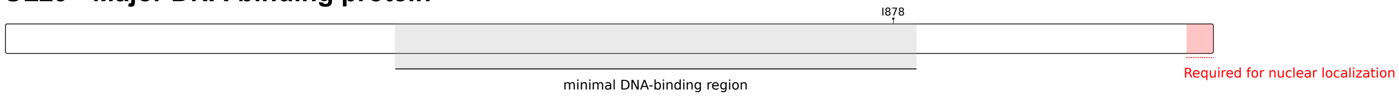


### UL36 - Large tegument protein

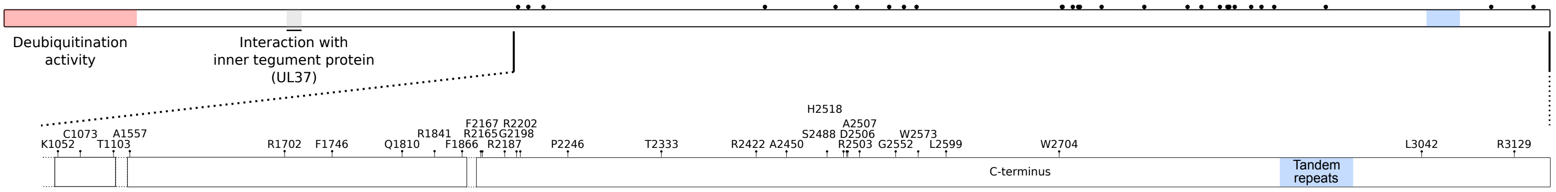


### UL55 - Nuclear protein UL55

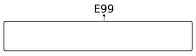


**Figure S1. Positive selection in UL26, UL29, UL36 and UL55.** Positively selected sites and functional domains were mapped onto HSV-1 proteins, as in Figure 2 and 3. For UL36, given the extended length of the protein sequence, positively selected sites were reported in the enlargement below.

**Supplementary Table S1. List of viral genome sequences.**

| Strain Name | Accession ID | Virus name (Common name; Strain abbreviation) | Virus Species | Host | Country |
|---|---|---|---|---|---|
| 2016-1040 | MG999886 | *Human herpesvirus 1* (Herpes simplex virus 1; HSV-1 2016-1040) | *Human alphaherpesvirus 1* | *Homo Sapiens* | USA |
| HSV-H12113 | MH999842 | *Human herpesvirus 1* (Herpes simplex virus 1; HSV-1 H12113) | *Human alphaherpesvirus 1* | *Homo Sapiens* | Finland |
| HSV-H1412 | MH999851 | *Human herpesvirus 1* (Herpes simplex virus 1; HSV-1 H1412) | *Human alphaherpesvirus 1* | *Homo Sapiens* | Finland |
| 172/2010 | LT594105 | *Human herpesvirus 1* (Herpes simplex virus 1; HSV-1 172/2010) | *Human alphaherpesvirus 1* | *Homo Sapiens* | Germany |
| HSV-1/0116209/India/2011 | KJ847330 | *Human herpesvirus 1* (Herpes simplex virus 1; HSV-1/0116209/India/2011) | *Human alphaherpesvirus 1* | *Homo Sapiens* | India |
| CJ394 | JN420340 | *Human herpesvirus 1* (Herpes simplex virus 1; HSV-1 CJ394) | *Human alphaherpesvirus 1* | *Homo Sapiens* | USA |
| CR38 | HM585508 | *Human herpesvirus 1* (Herpes simplex virus 1; HSV-1 CR38) | *Human alphaherpesvirus 1* | *Homo Sapiens* | China |
| E06 | HM585496 | *Human herpesvirus 1* (Herpes simplex virus 1; HSV-1 E06) | *Human alphaherpesvirus 1* | *Homo Sapiens* | Kenya |
| E07 | HM585497 | *Human herpesvirus 1* (Herpes simplex virus 1; HSV-1 E07) | *Human alphaherpesvirus 1* | *Homo Sapiens* | Kenya |
| E08 | HM585498 | *Human herpesvirus 1* (Herpes simplex virus 1; HSV-1 E08) | *Human alphaherpesvirus 1* | *Homo Sapiens* | Kenya |
| E13 | HM585502 | *Human herpesvirus 1* (Herpes simplex virus 1; HSV-1 E13) | *Human alphaherpesvirus 1* | *Homo Sapiens* | Kenya |
| H166 | KM222726 | *Human herpesvirus 1* (Herpes simplex virus 1; HSV-1 H166) | *Human alphaherpesvirus 1* | *Homo Sapiens* | USA |
| R11 | HM585514 | *Human herpesvirus 1* (Herpes simplex virus 1; HSV-1 R11) | *Human alphaherpesvirus 1* | *Homo Sapiens* | South Korea |
| S23 | HM585512 | *Human herpesvirus 1* (Herpes simplex virus 1; HSV-1 S23) | *Human alphaherpesvirus 1* | *Homo Sapiens* | Japan |
| CJ360 * | JN420339 | *Human herpesvirus 1* (Herpes simplex virus 1; HSV-1 CJ360) | *Human alphaherpesvirus 1* | *Homo Sapiens* | USA |
| 2003-16029 | KX574903 | *Human herpesvirus 2* (Herpes simplex virus 2; HSV-2 2003-16029) | *Human alphaherpesvirus 2* | *Homo Sapiens* | USA |
| 2004-4373 | KX574861 | *Human herpesvirus 2* (Herpes simplex virus 2; HSV-2 2004-4373) | *Human alphaherpesvirus 2* | *Homo Sapiens* | Peru |
| 2006-30937 | KX574866 | *Human herpesvirus 2* (Herpes simplex virus 2; HSV-2 2006-30937) | *Human alphaherpesvirus 2* | *Homo Sapiens* | Peru |
| 2006-49895 | KX574868 | *Human herpesvirus 2* (Herpes simplex virus 2; HSV-2 2006-49895) | *Human alphaherpesvirus 2* | *Homo Sapiens* | USA |
| 2007-38120 | KX574871 | *Human herpesvirus 2* (Herpes simplex virus 2; HSV-2 2007-38120) | *Human alphaherpesvirus 2* | *Homo Sapiens* | Zimbawe |
| 2007-38627 | KX574874 | *Human herpesvirus 2* (Herpes simplex virus 2; HSV-2 2007-38627) | *Human alphaherpesvirus 2* | *Homo Sapiens* | Peru |
| 2007-39712 * | KX574880 | *Human herpesvirus 2* (Herpes simplex virus 2; HSV-2 2007-39712) | *Human alphaherpesvirus 2* | *Homo Sapiens* | South africa |
| 2009-13458 | KX574889 | *Human herpesvirus 2* (Herpes simplex virus 2; HSV-2 2009-13458) | *Human alphaherpesvirus 2* | *Homo Sapiens* | USA |
| 2009-3532 | KX574892 | *Human herpesvirus 2* (Herpes simplex virus 2; HSV-2 2009-3532) | *Human alphaherpesvirus 2* | *Homo Sapiens* | South africa |
| 2009-4556 | KX574894 | *Human herpesvirus 2* (Herpes simplex virus 2; HSV-2 2009-4556) | *Human alphaherpesvirus 2* | *Homo Sapiens* | Kenya |
| 2011-21761 | KX574897 | *Human herpesvirus 2* (Herpes simplex virus 2; HSV-2 2011-21761) | *Human alphaherpesvirus 2* | *Homo Sapiens* | USA |
| HSV2-H1227 | KY922721 | *Human herpesvirus 2* (Herpes simplex virus 2; HSV-2 H1227) | *Human alphaherpesvirus 2* | *Homo Sapiens* | Finland |
| HSV2-H1229 | KY922722 | *Human herpesvirus 2* (Herpes simplex virus 2; HSV-2 H1229) | *Human alphaherpesvirus 2* | *Homo Sapiens* | Finland |
| 2008_15116 | MF621257 | *Human herpesvirus 2* (Herpes simplex virus 2; HSV-2 2008_15116) | *Human alphaherpesvirus 2* | *Homo Sapiens* | Kenya |
| CNS15 | MK105998 | *Human herpesvirus 2* (Herpes simplex virus 2; HSV-2 CNS15) | *Human alphaherpesvirus 2* | *Homo Sapiens* | USA |
| 105640 * | NC_023677 | Chimpanzee alpha-1 herpesvirus (Chimpanzee herpesvirus; ChHV 105640) | *Panine alphaherpesvirus 3* | *Pan troglodytes* | USA |
| 12930 | KY628971 | Macacine herpesvirus 1 (B virus; McHV-1 12930) | *Macacine alphaherpesvirus 1* | *Macaca mulatta* | USA |
| 1504-11 | KY628969 | Macacine herpesvirus 1 (B virus; McHV-1 1504-11) | *Macacine alphaherpesvirus 1* | *Macaca nemestrina* | USA |
| 16293 | KY628972 | Macacine herpesvirus 1 (B virus; McHV-1 16293) | *Macacine alphaherpesvirus 1* | *Macaca mulatta* | USA |
| 20620 | KY628973 | Macacine herpesvirus 1 (B virus; McHV-1 20620) | *Macacine alphaherpesvirus 1* | *Macaca mulatta* | USA |
| 24105-G | KY628974 | Macacine herpesvirus 1 (B virus; McHV-1 24105-G) | *Macacine alphaherpesvirus 1* | *Macaca mulatta* | USA |
| 26896-G | KY628975 | Macacine herpesvirus 1 (B virus; McHV-1 26896-G) | *Macacine alphaherpesvirus 1* | *Macaca mulatta* | USA |
| 31618-G | KY628978 | Macacine herpesvirus 1 (B virus; McHV-1 31618-G) | *Macacine alphaherpesvirus 1* | *Macaca mulatta* | USA |
| 32425-G | KY628981 | Macacine herpesvirus 1 (B virus; McHV-1 32425-G) | *Macacine alphaherpesvirus 1* | *Macaca mulatta* | USA |
| 7709642 | KY628982 | Macacine herpesvirus 1 | *Macacine alphaherpesvirus 1* | *Macaca silenus* | USA |

| | | | | | |
|---|---|---|---|---|---|
| | | (B virus; McHV-1 7709642) | | | |
| 8100812 | KY628968 | Macacine herpesvirus 1 (B virus; McHV-1 8100812) | *Macacine alphaherpesvirus 1* | *Macaca silenus* | USA |
| 9400371 | KY628983 | Macacine herpesvirus 1 (B virus; McHV-1 9400371) | *Macacine alphaherpesvirus 1* | *Macaca mulatta* | USA |
| E90-136 | KJ566591 | Macacine herpesvirus 1 (B virus; McHV-1 E90-136) | *Macacine alphaherpesvirus 1* | *Macaca fascicularis* | USA |
| KQ | KY628970 | Macacine herpesvirus 1 (B virus; McHV-1 KQ) | *Macacine alphaherpesvirus 1* | *Macaca nemestrina* | USA |
| M12-O | KY628985 | Macacine herpesvirus 1 (B virus; McHV-1 M12-O) | *Macacine alphaherpesvirus 1* | *Macaca radiata* | USA |
| E2490 ***** | NC_004812 | Macacine herpesvirus 1 (B virus; McHV-1 E2490) | *Macacine alphaherpesvirus 1* | *Macaca mulatta* | USA |
| A189164 | KF908239 | *Papiine herpesvirus 2* (Herpesvirus papio 2, PaHV-2 A189164) | *Papiine alphaherpesvirus 2* | - | USA |
| A951 | KF908242 | *Papiine herpesvirus 2* (Herpesvirus papio 2; PaHV-2 A951) | *Papiine alphaherpesvirus 2* | - | USA |
| OU2-5 | KF908241 | *Papiine herpesvirus 2* (Herpesvirus papio 2; PaHV-2 OU2-5) | *Papiine alphaherpesvirus 2* | *Papio anubis* | USA |
| OU4-2 | KF908244 | *Papiine herpesvirus 2* (Herpesvirus papio 2; PaHV-2 OU4-2) | *Papiine alphaherpesvirus 2* | *Papio ursinus* | USA |
| OU4-8 | KF908243 | *Papiine herpesvirus 2* (Herpesvirus papio 2; PaHV-2 OU4-8) | *Papiine alphaherpesvirus 2* | *Papio ursinus* | USA |
| X313 ***** | NC_007653 | *Papiine herpesvirus 2* (Herpesvirus papio 2; PaHV-2 X313) | *Papiine alphaherpesvirus 2* | *Papio anubis* | USA |
| SA8 ***** | NC_006560 | *Cercopithecine herpesvirus 2* (Simian agent 8; CeHV-2 SA8) | *Cercopithecine alphaherpesvirus 2* | *Cercopithecus aethiops* | - |

**Note: *** strains used for SLAC analysis.

**Supplementary Table S2. Herpes simplex genes excluded from the branch-sites analysis.**

| Gene Name | Motivation |
|---|---|
| *UL15* | Overlapping ORFs |
| *UL16* | Overlapping ORFs |
| *UL17* | Overlapping ORFs |
| *UL26.5* | Overlapping ORFs |
| *US4* | Excessive divergence and length among sequences |
| *US11* | Overlapping ORFs |
| *RL1* | Duplicated genes/no reliable orthologs |
| *RL2* | Duplicated genes/no reliable orthologs |
| *RS1* | Duplicated genes/no reliable orthologs |

**Supplementary Table S3. List of analyzed genes and dN/dS values**

| Gene Symbol | Protein Product Name | dN/dS |
|---|---|---|
| *UL1 ** | Envelope glycoprotein L | 0.219 |
| *UL2 ** | Uracil-DNA glycosylase | 0.106 |
| *UL3* | Nuclear protein UL3 | 0.174 |
| *UL5 ** | DNA replication helicase | 0.0876 |
| *UL4* | Nuclear protein UL4 | 0.182 |
| *UL6 ** | Capsid portal protein | 0.144 |
| *UL7 ** | Cytoplasmic envelopment protein 1 | 0.188 |
| *UL9 ** | DNA replication origin-binding protein | 0.129 |
| *UL8 ** | DNA helicase/primase complex-associated protein | 0.226 |
| *UL10 ** | Envelope glycoprotein M | 0.196 |
| *UL12 ** | Alkaline nuclease | 0.206 |
| *UL11 ** | Cytoplasmic envelopment protein 3 | 0.267 |
| *UL14 ** | Tegument protein UL14 | 0.242 |
| *UL13* | Serine/threonine-protein kinase UL13 | 0.172 |
| *UL18 ** | Triplex capsid protein 2 | 0.0933 |
| *UL20* | Envelope protein UL20 | 0.162 |
| *UL19** | Major capsid protein | 0.0827 |
| *UL21 ** | Tegument protein UL21 | 0.15 |
| *UL22 ** | Envelope glycoprotein H | 0.195 |
| *UL23 ** | Thymidine kinase | 0.234 |
| *UL24* | Nuclear protein UL24 | 0.153 |
| *UL25 ** | Capsid vertex component 2 | 0.107 |
| *UL26** | Capsid scaffolding protein | 0.196 |
| *UL28 ** | Tripartite terminase subunit 1 | 0.1 |
| *UL27** | Envelope glycoprotein B | 0.125 |
| *UL29** | Major DNA-binding protein | 0.0987 |
| *UL30** | DNA polymerase catalytic subunit | 0.116 |
| *UL32 ** | DNA packaging protein UL32 | 0.16 |
| *UL31 ** | Nuclear egress protein 1 | 0.133 |
| *UL33 ** | Tripartite terminase subunit 2 | 0.23 |
| *UL34 ** | Nuclear egress protein 2 | 0.14 |
| *UL35 ** | Small capsomere-interacting protein | 0.167 |
| *UL36** | Large tegument protein deneddylase | 0.229 |
| *UL37 ** | Inner tegument protein | 0.142 |
| *UL38 ** | Triplex capsid protein 1 | 0.147 |
| *UL39** | Ribonucleoside-diphosphate reductase large subunit | 0.184 |
| *UL40 ** | Ribonucleoside-diphosphate reductase small subunit | 0.111 |
| *UL41* | Virion host shutoff protein | 0.118 |
| *UL42 ** | DNA polymerase processivity factor | 0.216 |
| *UL43* | Membrane protein UL43 | 0.297 |
| *UL44* | Envelope glycoprotein C | 0.274 |
| *UL45* | Envelope protein UL45 | 0.189 |
| *UL46* | Tegument protein VP11/12 | 0.244 |
| *UL47* | Tegument protein UL47 | 0.203 |
| *UL48* | Tegument protein VP16 | 0.132 |
| *UL49A ** | Envelope glycoprotein N | 0.388 |
| *UL49* | Tegument protein VP22 | 0.123 |

| | | |
|---|---|---|
| *UL50 ** | Deoxyuridine 5'-triphosphate nucleotidohydrolase | 0.207 |
| *UL51 ** | Tegument protein UL51 | 0.124 |
| *UL52 ** | DNA primase | 0.135 |
| *UL53* | Envelope glycoprotein K | 0.141 |
| *UL54 ** | mRNA export factor (ICP27) | 0.223 |
| *UL55* | Tegument protein UL55 | 0.166 |
| *UL56* | Membrane protein UL56 | 0.21 |
| *US1* | Transcriptional regulator ICP22 | 0.286 |
| *US2* | Protein US2 | 0.172 |
| *US3* | Serine/threonine protein kinase US3 | 0.193 |
| *US5* | Envelope glycoprotein J | 0.458 |
| *US6* | Envelope glycoprotein D | 0.222 |
| *US7* | Envelope glycoprotein I | 0.293 |
| *US8* | Envelope glycoprotein E | 0.245 |
| *US8A* | Membrane protein US8.5 | 0.329 |
| *US9* | Envelope protein US9 | 0.147 |
| *US12* | TAP transporter inhibitor ICP47 | 0.301 |
| *US10* | Virion protein US10 | 0.273 |

**Note:** * *core* gene

**Supplementary Table S4. Likelihood ratio test (LRT) statistics for models of variable selective pressure on the Hominin-infecting SVs branch.**

| Gene | Alignment length (nt) | Tree length [a] | -2ΔlnL [b] | MA vs MA1 FDR corrected *p* value | sites BEB/MEME[c] |
|---|---|---|---|---|---|
| *UL1* | 774 | 5.2062 | 1.0366 | 0.48976 | |
| *UL2* | 1038 | 3.9312 | 1.1759 | 0.48126 | |
| *UL3* | 744 | 3.7135 | 1.1287 | 0.48126 | |
| *UL5* | 2715 | 2.4367 | 0.2981 | 0.77654 | |
| *UL4* | 618 | 4.1948 | 0 | 1.00000 | |
| *UL6* | 2091 | 2.8123 | 0.3888 | 0.73995 | |
| *UL7* | 894 | 3.4631 | 0.0042 | 0.97505 | |
| *UL9* | 2691 | 2.1922 | 1.0959 | 0.48126 | |
| *UL8* | 2391 | 3.8971 | 5.6139 | 0.08266 | |
| *UL10* | 1467 | 3.2181 | 16.3975 | **0.00187** | S51, R56, P58, A121, R126, Q163, L167, N238, N246, L269, C276, C300, R350, M357, L434, P445, R472 |
| *UL12* | 2100 | 2.6825 | 0 | 1.00000 | |
| *UL11* | 303 | 4.4465 | 0 | 1.00000 | |
| *UL14* | 663 | 1.8038 | 0 | 1.00000 | |
| *UL13* | 1569 | 2.8092 | 0.1301 | 0.85964 | |
| *UL18* | 954 | 2.2424 | 0.1742 | 0.82297 | |
| *UL20* | 681 | 3.0543 | 1.4337 | 0.41900 | |
| *UL19 Reg 1* | 2208 | 2.0752 | 0.7785 | 0.58649 | |
| *UL19 Reg 2* | 1911 | 1.9389 | 3.5890 | 0.18116 | |
| *UL21* | 1626 | 3.4383 | 2.7415 | 0.23791 | |
| *UL22* | 2517 | 3.6050 | 10.6183 | **0.01635** | Y85, F163, E170, F174, H185, G188, R190, R195, R225, R233, E304, P317, E347, N473, L496, A532, W581, Y637 |
| *UL23* | 1137 | 3.1730 | 3.3122 | 0.20080 | |
| *UL24* | 882 | 2.6549 | 0.2422 | 0.78595 | |
| *UL25* | 1770 | 2.3141 | 2.9889 | 0.22983 | |
| *UL26 Reg 1* | 834 | 2.1814 | 4.1068 | 0.16410 | |
| *UL26 Reg 2* | 1281 | 3.5030 | 8.2015 | **0.03395** | Y374, A379, S612 |
| *UL28* | 2451 | 1.6837 | 1.7080 | 0.37733 | |
| *UL27 Reg 2* | 2364 | 1.6158 | 7.2564 | **0.04298** | R136, A334 |
| *UL29 Reg 1* | 2055 | 2.2775 | 0.2397 | 0.78595 | |
| *UL29 Reg 2* | 1557 | 1.9996 | 9.7336 | **0.02201** | I878 |
| *UL30 Reg 1* | 2805 | 2.2478 | 2.0888 | 0.30949 | |
| *UL30 Reg 2* | 1026 | 1.6936 | 1.5944 | 0.38691 | |
| *UL32* | 1833 | 2.2573 | 2.3234 | 0.28191 | |
| *UL31* | 933 | 1.8317 | 0.0772 | 0.91968 | |
| *UL33* | 411 | 1.8974 | 0 | 1.00000 | |
| *UL34* | 861 | 2.8715 | 1.0891 | 0.48126 | |
| *UL35* | 345 | 3.9459 | 0 | 1.00000 | |
| *UL36 Reg 1* | 2793 | 3.4117 | 0.4144 | 0.73995 | |
| *UL36 Reg 2* | 1521 | 2.4680 | 8.9815 | **0.02489** | K1052, C1073, T1103 |
| *UL36 Reg 3* | 4689 | 2.7863 | 28.9285 | **0.00001** | A1557, R1702, F1746, Q1810, R1841, F1866, R2165, F2167, R2187, G2198, R2202, P2246, T2333, R2422, A2450, S2488, R2503, D2506, A2507, H2518, G2552, W2573, L2599, W2704, L3042, R3129 |
| *UL37* | 3372 | 2.7079 | 1.9638 | 0.32669 | |
| *UL38* | 1407 | 2.9530 | 0.2442 | 0.78595 | |
| *UL39 Reg 1* | 2997 | 2.9086 | 1.4084 | 0.41900 | |
| *UL39 Reg 2* | 822 | 1.6584 | 2.8690 | 0.22983 | |
| *UL40* | 1050 | 1.8324 | 0.9592 | 1.00000 | |
| *UL41* | 1506 | 2.8685 | 0.3180 | 0.77438 | |
| *UL42* | 1614 | 3.4912 | 0 | 1.00000 | |
| *UL43* | 1353 | 4.0241 | 0.7059 | 0.60956 | |
| *UL44* | 1365 | 3.9588 | 6.6507 | 0.05032 | |

| | | | | | |
|---|---|---|---|---|---|
| *UL45* | 522 | 3.8573 | 6.4226 | 0.05875 | |
| *UL46* | 2376 | 2.9204 | 7.9589 | **0.03493** | E32, E265, T311, I421, A535, W629, Y649 |
| *UL47 Reg 1* | 858 | 4.2778 | 0.3595 | 0.75589 | |
| *UL47 Reg 2* | 1377 | 2.8867 | 4.8996 | 0.10894 | |
| *UL48* | 1491 | 3.2001 | 3.9319 | 0.17293 | |
| *UL49A* | 279 | 6.2218 | 3.7508 | 0.18116 | |
| *UL49* | 963 | 4.9452 | 4.5255 | 0.09128 | |
| *UL50* | 1164 | 4.3078 | 1.6095 | 0.38691 | |
| *UL51* | 771 | 2.4518 | 3.5497 | 0.18116 | |
| *UL52* | 3258 | 2.8502 | 0.1823 | 0.82297 | |
| *UL53* | 1014 | 3.8113 | 2.9231 | 0.22983 | |
| *UL54* | 1683 | 3.2669 | 0.6472 | 0.62738 | |
| *UL55* | 570 | 3.9219 | 9.2010 | **0.02489** | E99 |
| *UL56* | 828 | 6.4393 | 3.6732 | 0.18116 | |
| *US1* | 1386 | 5.3915 | 14.8083 | **0.00290** | T47, M90, S91, M147, C231, S247, T250, C274, C278 |
| *US2* | 954 | 4.2054 | 2.4006 | 0.27670 | |
| *US3* | 1467 | 3.0700 | 2.6557 | 0.24297 | |
| *US5* | 426 | 6.9588 | 0 | 1.00000 | |
| *US6* | 1200 | 3.5028 | 5.5849 | 0.08266 | |
| *US7* | 1338 | 4.4700 | 2.8513 | 0.22983 | |
| *US8* | 1500 | 4.3746 | 7.2839 | **0.04298** | A5, F9, C18, P40, A98, E227, T269, G313, H328, P351, Q352, A411, L429, L433, R446, G481, E493 |
| *US8A* | 345 | 5.1603 | 0.0000 | 1.00000 | |
| *US9* | 282 | 4.3681 | 2.2236 | 0.29181 | |
| *US10* | 765 | 4.5635 | 0.4088 | 0.73995 | |
| *US12* | 333 | 8.6430 | 10.9987 | **0.01635** | A4, D27, K31, R32, R34, R41 |

**Notes:**

[a] Branch length is defined as number of nucleotide substitutions per codon

[b] 2ΔlnL: twice the difference of the natural logs of the maximum likelihood of the models being compared

[c] Positions refer to proteins of HHV-1 Strain 17 (NC_001806).
